# Early altered directionality of resting brain network state transitions in the TgF344-AD rat model of Alzheimer’s disease

**DOI:** 10.1101/2024.02.05.578351

**Authors:** Sam De Waegenaere, Monica van den Berg, Georgios A. Keliris, Mohit H. Adhikari, Marleen Verhoye

**Author notes:** co-senior authors (Mohit H. Adhikari, Marleen Verhoye).

## Abstract

Alzheimer’s disease (AD) is a progressive neurodegenerative disease resulting in memory loss and cognitive decline. Synaptic dysfunction is an early hallmark of the disease whose effects on whole-brain functional architecture can be identified using resting-state functional MRI (rsfMRI). Insights into mechanisms of early, whole-brain network alterations can help our understanding of the functional impact of AD’s pathophysiology. Here, we obtained rsfMRI data in the TgF344-AD rat model at the pre- and early-plaque stages. This model recapitulates the major pathological and behavioural hallmarks of AD. We used co-activation pattern (CAP) analysis to investigate if and how the dynamic organization of intrinsic brain functional networks states, undetectable by earlier methods, is altered at these early stages. We identified and characterized six intrinsic brain states as CAPs, their spatial and temporal features, and the transitions between the different states. At the pre-plaque stage, the TgF344-AD rats showed reduced co-activation of hub regions in the CAPs corresponding to the default mode-like and lateral cortical network. Default mode-like network activity segregated into two distinct brain states, with one state characterised by high co-activation of the basal forebrain. This basal forebrain co-activation was reduced in TgF344-AD animals mainly at the pre-plaque stage. Brain state transition probabilities were altered at the pre-plaque stage between states involving the default mode-like network, lateral cortical network, and basal forebrain regions. Additionally, while the directionality preference in the network-state transitions observed in the wild-type animals at the pre-plaque stage had diminished at the early-plaque stage, TgF344-AD animals continued to show directionality preference at both stages. Our study enhances the understanding of intrinsic brain state dynamics and how they are impacted at the early stages of AD, providing a nuanced characterization of the early, functional impact of the disease’s neurodegenerative process.

## Introduction

Resting-state functional MRI (rsfMRI), offering insight into the brain’s functional organisation and connectivity, has emerged as a powerful non-invasive tool in Alzheimer’s Disease (AD) research. Insights obtained using this imaging technique have shown potential for early detection of AD, as it identified brain network alterations and abnormalities before the onset of clinical signs (Sheline and Raichle, 2013). RsfMRI records spontaneous low-frequency fluctuations in the blood-oxygenation-level-dependent (BOLD) signals as an indirect measure of neuronal activity at rest, such as during quiet wakefulness or while lying still with eyes closed. It allows for the mapping of brain-wide functional connectivity (FC), defined as the strength of the temporal correlation of BOLD activity between spatially distant regions, and in turn unveils resting-state networks (RSNs) constituted by brain regions with correlated activity. Predominantly active during rest and associated with introspection and memory, the Default Mode Network (DMN) is a particularly relevant RSN in AD (Badhwar et al., 2017), as its reduced connectivity is indicative of the disease’s progression (Alves et al., 2019, van den Heuvel and Sporns, 2013, Greicius et al., 2004, Ramzan et al., 2019). Furthermore, the DMN’s anti-correlation with the Task-Positive Network (TPN), which is engaged during cognitive tasks, is diminished in AD, thus reducing the segregation between DMN and TPN (Belloy et al., 2018b). Understanding changes in FC within and between the DMN, TPN, and other RSNs has shown to be highly relevant to elucidate AD’s impact on brain function and potential for early detection and monitoring of disease progression (Ibrahim et al., 2021, Agosta et al., 2012).

Animal models of AD provide critical insights into the disease’s pathophysiology and enable the exploration of especially early impact of AD pathology. The short lifespan allows for monitoring the evolution of the disease phenotype in a relatively brief period, even before onset of the behavioural phenotype. One such model, the TgF344-AD rat, is characterized by the presence of human APP_swe_ and PS1_ΔE9_ mutations, leading to the progressive build-up of amyloid beta (Aβ) deposits starting at 6 months of age (Cohen et al., 2013, Joo et al., 2017), accompanied by accumulation of phosphorylated tau (pTau) (Joo et al., 2017, Rorabaugh et al., 2017), gliosis, neuronal loss and cognitive impairments similar to those observed in human AD (Braak et al., 2011, Munoz-Moreno et al., 2018, Cohen et al., 2013). Impairments in spatial reference learning, spatial navigation and working memory have been observed starting at 6 months of age (Tournier et al., 2021, Berkowitz et al., 2018), with anxiety-like behaviour preceding already at the age of 4 months (Pentkowski et al., 2018).

RsfMRI in rodents shows similar features as in humans; in particular analogues of RSNs have been identified in mice and rats (Lu et al., 2012). In previous work using different MRI modalities including rsfMRI in TgF344-AD female rats (Anckaerts et al., 2019) we have shown that changes in FC preceded structural changes. A small decrease in FC at the age of 6 months (in connections involving the hippocampus, cingulate, retrosplenial and sensorimotor cortices) was followed by widespread hypoconnectivity at 10 months. Another study (Tudela et al., 2019) showed lowered DMN overall mean connectivity at 5 months and decreased anterior-posterior DMN connectivity at 15 months. More evidence of network-level alterations preceding plaque formation was found in regional FC alterations at 5 months, located in the right insular cortex, amygdala and hypothalamus (Munoz-Moreno et al., 2018, Munoz-Moreno et al., 2020). The combined findings of these studies emphasize the relevance of this model in characterizing not only AD’s pathology and behavioural phenotype, but also its impact, especially at early stages, on brain function.

These rsfMRI studies assumed static, constant FC throughout the scan. However, advances like sliding window FC analyses have revealed FC fluctuations that are missed with static FC (Hutchison et al., 2013, Calhoun et al., 2014), and changes in activity patterns over the scan’s duration (Xu et al., 2022, Lurie et al., 2020, Maltbie et al., 2022). One such approach uncovers the quasi-periodic patterns (QPPs) (Majeed et al., 2011), which are recurring spatiotemporal patterns of brain activity within a predefined time window. QPPs are obtained using a sliding window approach, where repetitive patterns of brain-wide neural activity are detected and averaged across them. Depending on the window size, QPPs can range from a single spatiotemporal brain state pattern to a sequence of them. In contrast to sliding window FC approaches, which yield one FC matrix per window, QPPs facilitate further insights into the dynamics of the activity states within the window itself (Yousefi and Keilholz, 2021, Yousefi et al., 2018, Belloy et al., 2021, Belloy et al., 2018a). In the context of AD, QPP’s have been used to detect early alterations in preclinical models (van den Berg et al., 2022, Belloy et al., 2018b). Our study using QPPs in TgF344-AD rats (van den Berg et al., 2022) highlighted a specific alteration: while overall FC between the basal forebrain (BFB) and the default mode-like network (DMLN) showed no notable differences, there was a distinct reduction in their coactivation within the QPP in TgF344-AD rats already at the pre-plaque stage of 4 months of age. This finding underscores the nuanced, early changes in neural interactions and disease processes in AD, detectable through dynamic QPP analysis but overlooked by conventional FC methods. However, insights gained from QPPs are inherently limited to the recurring states, covering only parts of entire scan. Additionally, an arbitrary selection of window size for analysis can yield varying outcomes, potentially missing dynamics that occur outside or across the chosen windows.

Co-activation patterns (CAPs) have been proposed as an alternative framework for investigating the brain’s dynamic states. CAPs are based on point process models (Tagliazucchi et al., 2012) and use clustering (Liu and Duyn, 2013) to divide all time frames within a scan, based on spatial similarity of their voxel-level BOLD signals, into a few (typically 6-10) functional coactivation states. They offer several advantages over QPPs, including the ability to capture and quantify inter-CAP transitions and higher statistical power due to single-frame resolution. Utilizing CAPs as a novel analytical tool has shown promise in classifying and differentiating properties associated with various physiological and pathological conditions (Adhikari et al., 2020, Adhikari et al., 2023, Gutierrez-Barragan et al., 2019, Gutierrez-Barragan et al., 2022, Liu et al., 2018, Vasilkovska et al., 2023, An et al., 2024, Lee et al., 2023, Li et al., 2021). Recent studies employing CAP analysis in mouse models of neurodegenerative disorders (Adhikari et al., 2023, Gutierrez-Barragan et al., 2022) have demonstrated their predictive capacity in distinguishing model mice from the wildtype at different stages of the disease.

In this work, we performed a resting-state CAP analysis on our previously obtained longitudinal dataset in the TgF344-AD rat model of AD at the ages of 4 and 6 months, corresponding to the pre- and early-plaque stages of disease progression (van den Berg et al., 2022). Our first aim was to investigate the genotype and age effects on the spatial and temporal properties of CAPs. Then we assessed the capability of CAP metrics to accurately identify the genotypes, by investigating their classification capabilities with specific interest for the pre-plaque stage. Finally, and most importantly, we characterised inter-CAP transitions, to investigate if and how the dynamic organization of intrinsic brain functional networks states, undetectable by earlier methods, is altered at these early stages in this model of AD.

### Materials & Methods

The data utilized in this study were initially obtained and published in a previous manuscript (van den Berg et al., 2022). Acquisition and rsfMRI pre-processing procedures are reiterated here for completeness.

### Animals and study design

All TgF344-AD and F344 rats used for this study were bred in-house. Heterozygous TgF344-AD rats were obtained from the RRRC (RRID: RGD_10401208) and were crossed with F344 rats from Charles River, Italy. Only male offspring were included in this study. A total of 15 male TgF344-AD (TG) rats and 11 wildtype (WT) littermates underwent rsfMRI scans at 4 (4M) and 6 (6M) months of age. For the current analysis, we only included the animals with data at both ages (TG: n=13, WT: n=11). Animals were group housed under controlled conditions, with a 12-hour light/dark cycle (lights on at 7 am) and a temperature maintained at 20-24 °C with 40-60% humidity. They had access to standard food and water ad libitum. All procedures were performed in strict accordance with the guidelines approved by the European Ethics Committee (decree 2010/63/EU) and were approved by the Committee on Animal Care and Use at the University of Antwerp, Belgium (approval number: 2019-06).

### *In vivo* MRI procedures

The rats were anesthetized using isoflurane, with 5% concentration for induction and 2% during handling. They were intubated endotracheally and mechanically ventilated at a rate of 70 breaths per minute. The head was secured with ear bars, and a cannula was inserted into the tail vein for intravenous administration of medetomidine and pancuroniumbromide. Initial bolus injection of medetomidine (0.05 mg/kg, Domitor®, Pfizer, Germany) and pancuroniumbromide (0.5 mg/kg, VWR, Belgium) was followed by a continuous infusion of medetomidine (0.1 mg/kg/h) and pancuroniumbromide (0.5 mg/kg/h) starting 15 minutes after the bolus injection. The concentration of isoflurane was gradually reduced to 0.4%. Throughout the procedure, the animals’ physiological parameters were closely monitored. Body temperature was maintained at 37 ± 0.5 °C using a feedback-controlled warm air circuitry system (MR-compatible Small Animal Heating System, SA Instruments, Inc., USA). Breathing rate was recorded using a pressure-sensitive pad (MR-compatible Small Animal Monitoring and Gating system, SA Instruments, Inc., USA) and heart rate and blood oxygenation were monitored using a pulse oximeter placed on the hind paw (MR-compatible Small Animal Monitoring and Gating System, SA Instruments, Inc., USA).

### MRI Acquisition

Data were obtained using a 9.4 T Biospec MRI system (Bruker BioSpin, Germany) and a 2×2 array receiver head radiofrequency coil, using Paravision 6 software. T2-weighted TurboRARE images were acquired in three directions to ensure consistent slice positioning. Magnetic field inhomogeneity was corrected with local shimming within an ellipsoid volume of interest that covered the brain.

rsfMRI data were collected using a single-shot gradient echo EPI-sequence with the following parameters: repetition time (TR) 600 ms, echo time (TE) 18 ms, field of view (FOV) (30 × 30) mm², matrix size [96 × 96], 12 coronal slices of 0.9 mm thickness with a 0.1 mm gap, 1000 repetitions. rsfMRI scans started 35 minutes after initial bolus administration and lasted 10 minutes.

After completion of the scan, infusion of medetomidine and pancuroniumbromide anaesthesia was stopped, and the isoflurane level was increased to 2%. T2-weighted 3D images were then acquired using a 3D RARE sequence (TR 2500 ms, effective TE 44 ms, F OV (30 × 30 × 22) mm³, matrix size of [256 × 256 × 128], RARE-factor 16) for registration purposes. At the end of the session, the animals received a subcutaneous injection of 0.1 mg/kg atipamezole (Antisedan®, Pfizer, Germany) to counteract the effects of medetomidine anaesthesia. Animals were placed on a second ventilator and heating pad to facilitate recovery. Except two 4-month-old TgF344-AD rats who did not recover due to premature extubation, all animals recovered within one hour after the scan session.

### Pre-processing

Pre-processing of the data including debiasing, realignment, co-registration, normalization, and smoothing was performed using SPM12 software (Statistical Parametric Mapping). First, debiasing was applied to the 3D RARE scans to eliminate intensity gradients caused by the RF coil. A study-specific 3D template was created from a subset of debiased 3D RARE scans using Advanced Normalization Tools (ANTs). The rsfMRI images were then realigned to the first image using a 6-parameter rigid body spatial transformation estimated using a least-square approach. Then, the rsfMRI images were co-registered to the corresponding anatomical 3D scan of the same session using a global 12-parameter affine transformation with mutual information as the similarity metric. The anatomical 3D scan was normalized to the study-specific 3D template using a global 12-parameter affine transformation followed by a non-linear deformation protocol. This combined transformation was applied to realign all EPI (echo-planar imaging) images and normalize them to the 3D RARE template. Additionally, ventricles were masked out of the data, and in-plane smoothing was performed using a Gaussian kernel with a full width at half maximum of twice the voxel size.

We used custom in-house MATLAB scripts for filtering, detrending, signal regression and z-scoring. To avoid transient effects, six time frames at the start and at the end of the scan were removed before the filtering process. The images then underwent voxel-wise filtering with a 0.01–0.2 Hz Butterworth band-pass filter, followed by another cut of four time frames at the start and end of the scan to minimise boundary effects of filtering. After quadratic detrending, the signal from white matter and ventricles (obtained using an atlas normalised to the template) was then regressed out from each voxel’s BOLD signal timetrace, which was subsequently normalized to have a zero mean and unit variance.

### CAP extraction

To analyse temporal fluctuations of neural activity, we employed an approach by Gutierrez-Barragan et al (Gutierrez-Barragan et al., 2019). Pre-processed rsfMRI data of all subjects (both WT and TG animals) at both ages (4M and 6M) were masked using an atlas-based whole brain mask excluding ventricles and concatenated into a single combined image-series. The image series was transformed into N m-dimensional vectors with N the total number of frames (across all subjects and ages) and m the total number of voxels within the whole-brain mask. We then applied a threshold so that, for each frame, z-scored BOLD signals of voxels within the top 10^th^ and bottom 5^th^ percentiles were kept while signals for all other voxels were set to zero. All time frames were then clustered according to spatial similarity, using K-means++ clustering algorithm. Specifically, the correlation distance (1 – Pearson’s correlation) between every pair of m-dimensional vectors was used to determine the spatial dissimilarity of the activation pattern of each frame with that of every other frame.

The K-means++ algorithm allocates the vectors into K clusters, minimising the sum of within-cluster distance *D* = 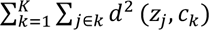. *K* represents the number of clusters, and *d*(*z*_*j*_, *c*_*k*_) refers to the correlation distance between the centroid *c*_*k*_ of the *k*^th^ cluster and *z*_*j*_, the z-scored BOLD signals of all voxels at the *j*^th^ time frame assigned to that cluster. To ensure an optimal selection of initial cluster centroids, the K-means++ algorithm favours distant centroids to be selected as initial seeds. We systematically explored a range of 2 to 30 ‘time frame’ clusters, calculating the amount of across-subject variance explained in each case using the following approach:

- Within-cluster variance, 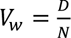, with *N* being the total number of time frames.
- Between cluster variance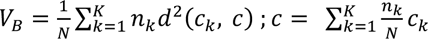. The sum of squared correlation distances between the global centroid, *c*, and each cluster centroid, *c*_*k*_, weighted by the number of frames in each cluster.
- Explained variance = 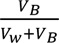. This is the ratio of between-cluster variance to the sum of between-cluster variance and within-cluster variance.

We then generated a plot showing the explained variance as a function of the number of clusters, ranging from 2 to 30. We identified the optimal number of clusters as the point at which the variance reached a saturation level, commonly known as the elbow point. Additionally, for each partition with k clusters, we calculated the fractional gain in explained variance compared to the partition with k-1 clusters. To ensure the validity of the elbow point, we confirmed that the fractional gain in explained variance for partitions with higher numbers of clusters remained below 0.5%. For each cluster, the original, whole brain, z-scored BOLD signals (without thresholding used for K-means clustering and calculation of explained variance) were averaged voxel-wise across all cluster time frames, resulting in combined cohort-level CAPs. Similarly, we performed averaging across the genotype-age-specific time frames within each cluster to obtain the CAPs for each subgroup (WT 4M, WT 6M, TG 4M, TG 6M).

### CAP analysis

We conducted a two-tailed, one-sample T-test (Bonferroni corrected, p<0.01) to identify significantly activated or deactivated voxels, for each CAP, using its occurrences in the combined image-series and its respective age-specific WT or TG sections. One-sample T-statistics for significant voxels were visualized as a brain-wide map for each CAP.

Next, we calculated the following measures of temporal and spatial properties of each CAP:

- Voxel-level activations: The mean activation observed in all occurrences of a CAP within the genotype-age-group-level image-series.
- Occurrence percentage: The percentage of time frames within a subject’s image-series that corresponded to a specific CAP.
- Duration: The average number of consecutive frames representing a CAP across all its occurrences within a subject.

### Statistical comparisons for spatial and temporal properties

To assess effects of age and genotype on voxel-wise spatial activation within each CAP, we used two-way ANOVA (age, genotype, age*genotype). Where the interaction effect between age and genotype did not reach statistical significance (p > 0.05), it was excluded prior to assessing main effects of age and genotype. For voxels that demonstrated significant interaction effect, post-hoc analyses (t-test) were performed, followed by corrections for multiple comparisons using Benjamini-Hochberg method to control for the false discovery rate (FDR).

Similarly, the effects of age, genotype, and their interaction on the temporal characteristics of CAPs were analysed using repeated measures 2-way ANOVA. In situations where no significant interaction effect was observed (p > 0.05), it was omitted from subsequent main effect analysis. The post hoc comparisons for occurrence, and duration underwent correction for multiple comparisons, using the Benjamini-Hochberg method to control the FDR.

### Classification

We used a multinomial logistic regression (MLR) model with regularization to categorize the subjects into their four distinct genotype-age-groups (WT 4M, WT 6M, TG 4M, TG 6M) by analysing both temporal (occurrence and duration) and spatial features (z-scored BOLD signal intensities of voxels, found to be significant in at least one of the four groups (p < 0.01, one-sample T-test, Bonferroni-corrected). The classifier’s training phase included 80% of the subjects, randomly selected in age and genotype stratified manner, to ensure balanced class representation. The remaining 20% of subjects were used as a validation set to calculate prediction accuracy. The splitting of the cohort into training and validation sets was iterated 50 times to yield a distribution of classification accuracy scores.

To establish a baseline for chance-level accuracy, we randomized the class labels within the same training-validation sets split for each of the 50 iterations and obtained the chance-level accuracy values. The median accuracy achieved through our classification was then statistically compared against this chance-level baseline via the Wilcoxon sign rank test, confirming whether our model’s predictions were significantly beyond what could be expected by random guessing.

To avoid a bias in the classifier where information of the validation group would cause more accurate prediction than otherwise possible, we adjusted the CAP extraction process to only include data from the training set when determining genotype-age-group-level CAPs. These subgroup-specific CAPs were then used as a reference to identify and calculate individual-specific CAP attributes within each subject’s scans.

By pooling the test-set results across all 50 iterations, we quantified the proportion of correctly classified subjects per class and those misclassified as one of the other three categories. The outcomes of this classification were compiled in a confusion matrix, where the prediction accuracy for each class is detailed on the diagonal, and the likelihood for misclassification between classes is presented in the off-diagonal cells. Furthermore, we verified if classification would be improved when predicting genotypes per age. Using the same methodology, we additionally performed two-class classification of the genotypes at each age.

### Inter-CAP Transition probabilities

We calculated the inter-CAP transition probabilities at the group-level at each age. We first calculated persistence probability of each CAP by identifying the total number of self-transitions (same CAP at t and t+1) from all subjects within the genotype-age-group and dividing them by the total number of transitions (self and non-self) made by that CAP. The transition probability, p_ij_, of CAP i to CAP j was calculated by dividing the total number of transitions between CAP i to CAP j by the total number of non-self-transitions (by excluding persistent or self-transitions) by CAP i to all other CAPs. The significance of transition probability between a CAP pair was tested by comparing it with a distribution of surrogate transitions probabilities for that pair. Surrogate transition probabilities were calculated using 10000 surrogate sequences of CAPs in each subject obtained by randomly permuting the CAP identities of time frames from that subject. The p-values for each CAP-pair were calculated by permutation testing, as the ratio of number of surrogates with transition probability greater than the observed value and the number of surrogates. Subsequently, p-values were FDR-corrected for all CAP pairs. We also calculated the difference in transition probability p_ij_ and p_ji_ in each genotype-age-group and assessed if the difference is statistically significant by comparing with a distribution of surrogate differences. The significance of this difference was tested for those CAP pairs which showed significant transition probability in at least one direction and all the p-values were FDR-corrected for the number of null hypotheses tested. Finally at each age, we calculated the inter-genotypic difference between transition probabilities for CAP pairs that were significant in at least one group, as well as between persistent probabilities for CAPs. Each such difference was statistically compared with a distribution of surrogate differences in a permutation test to assess its significance and the subsequent p-values were corrected for multiple comparisons using FDR.

## Results

### CAP analysis reveals six brain states

We initially determined the optimal number of CAPs. Figure 1A illustrates the variance explained across subjects through partitions, with an increasing number of clusters, of the concatenated image-series from all subjects (11 WT, 13 TG) at both ages. We identified 6 as the optimal number of clusters, with the explained variance showing a clear elbow point at 6 (Figure 1A) and a fractional gain in variance less than 0.5% (Figure 1B). We then generated CAPs by computing voxel-wise averages of z-scored BOLD signals across all time frames sharing identical cluster membership (Figure 1C).

**Figure 1.**
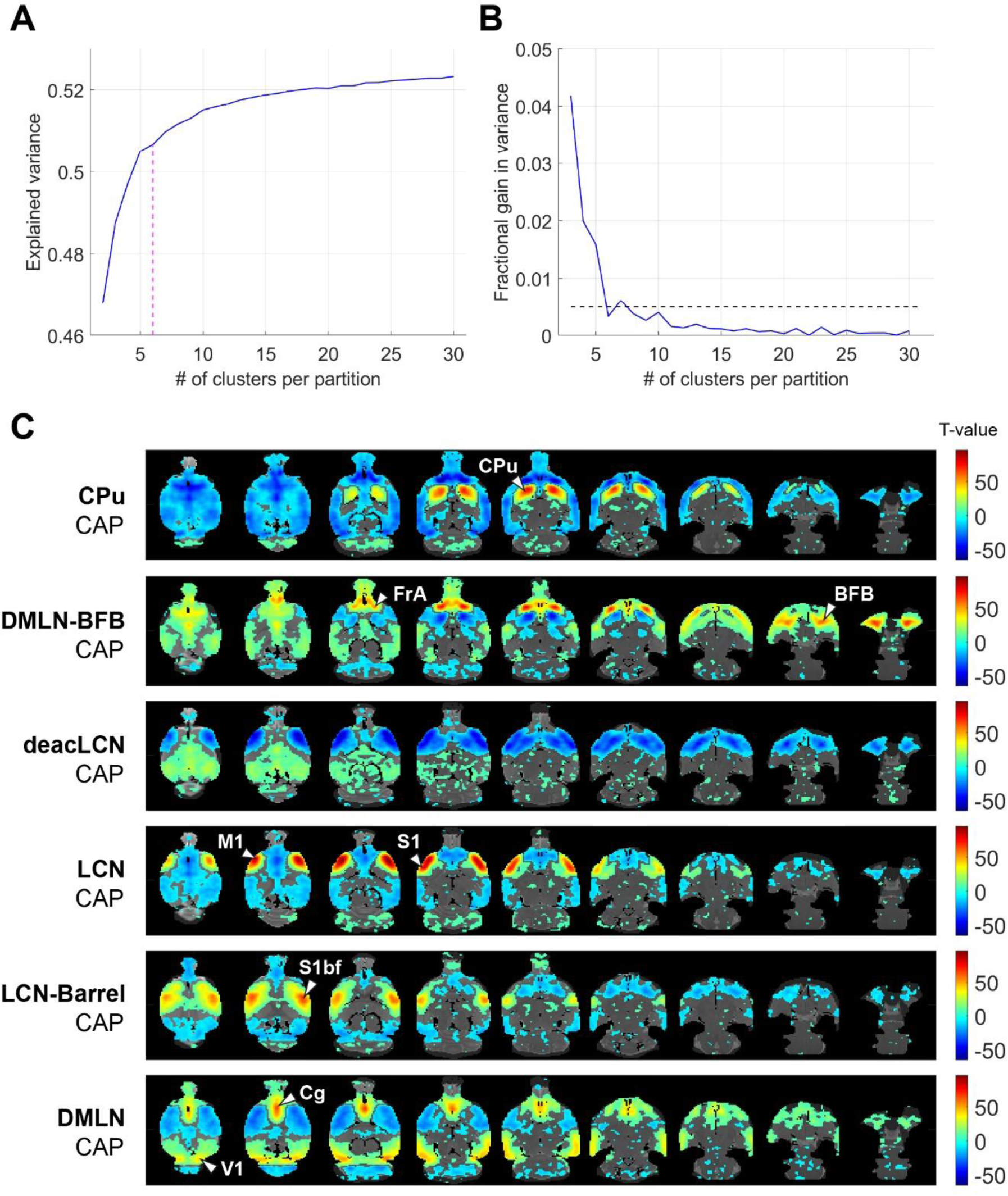
Selection of optimal CAP partition and visualisation of cohort-level CAPs. **(A)** Across-subject variance explained as a function of number of clusters in the partition of the combined image-series. Magenta dashed line indicates the elbow point beyond which the explained variance is found to saturate in each case. We found 6 clusters were sufficient to explain ∼50% of variance. **(B)** Fractional gain in the explained variance with k clusters versus k-1 clusters. The elbow point is the first k which has a fractional gain lower than 0.5% (shown by dashed line). **(C)** Cohort-level (across both genotypic groups and ages) one-sample T-statistic maps of the CAPs. Voxels with positive T-values (yellow to red) indicate a significant (p < 0.01; Bonferroni corrected) activation, i.e., higher BOLD signal averaged across the CAP-timeframes as compared to the BOLD signal averaged across all time frames. Voxels with negative T-values (cyan to blue) indicate significant deactivation, i.e., lower BOLD signal averaged across the CAP-timeframes as compared to the BOLD signal averaged across all time frames. CPu, caudate putamen; FrA, frontal association cortex; BFB, basal forebrain; M1, primary motor cortex; S1, secondary motor cortex; S1bf, barrel cortex; V1, primary visual cortex; Cg, cingulate cortex.

### CAPs are characterised by hubs of (in)activation spanning multiple known RSNs

The 6 identified CAPs were sorted into distinct CAP-anti CAP pairs, ordered first by occurrence per CAP then rearranged to form pairs sorted from strongest to weakest Pearson anti-correlation (Figure 1C). The CAPs were named based on prominent features in their spatial representations.

The *LCN-Barrel* CAP and *DMLN* CAP exhibited a spatial pattern very similar to the LCN and DMLN CAPs, described previously in a Huntington’s disease mice study (Adhikari et al., 2023), respectively. In the *LCN-Barrel* CAP, co-activation of lateral cortical network (LCN) regions such as the somatosensory, motor, insular cortices, and a main hub of activation in the barrel cortex was exhibited with simultaneous co-deactivation of default mode-like network (DMLN) regions such as cingulate, visual, retrosplenial and temporal association cortices. The *DMLN* CAP demonstrated an anti-correlated pattern, where DMLN regions were co-activated and LCN regions were co-deactivated.

In anti-correlated CAPs *CPu* and *DMLN-BFB*, co-activation and co-deactivation of the caudate putamen was paired with widespread cortical co-deactivation and co-activation, respectively. The *CPu* CAP exhibited a distinct and highly localised co-activation across the entire caudate putamen, while the *DMLN-BFB* CAP was mainly characterised by strong co-activation in the frontal association cortex and the BFB. In the *DMLN-BFB* CAP, the primary somatosensory cortex as well as the CPu also showed pronounced co-deactivation.

The *deacLCN* and *LCN* CAP formed an anti-correlated pair mainly characterised by motor and primary somatosensory regions. The *deacLCN* CAP exhibited localized co-deactivation in these regions, paired with widespread co-activation in central regions. In the *LCN* CAP, motor and primary somatosensory cortex formed hubs of highly localised strong co-activation, with concurrent widespread (sub)cortical de-activation.

### CAP activations show localised inter-genotypic differences mainly at 4 months

Next, we assessed the effects of genotype and/or age on voxel-level spatial activations in each of the six CAPs, using two-way ANOVA (age, genotype, age*genotype). Figure 2 and Figure 3 show the six CAPs in each genotypic group and the post-hoc comparisons of spatial activations between the two genotypes at 4M (Figure 2) and 6M (Figure 3) respectively.

**Figure 2.**
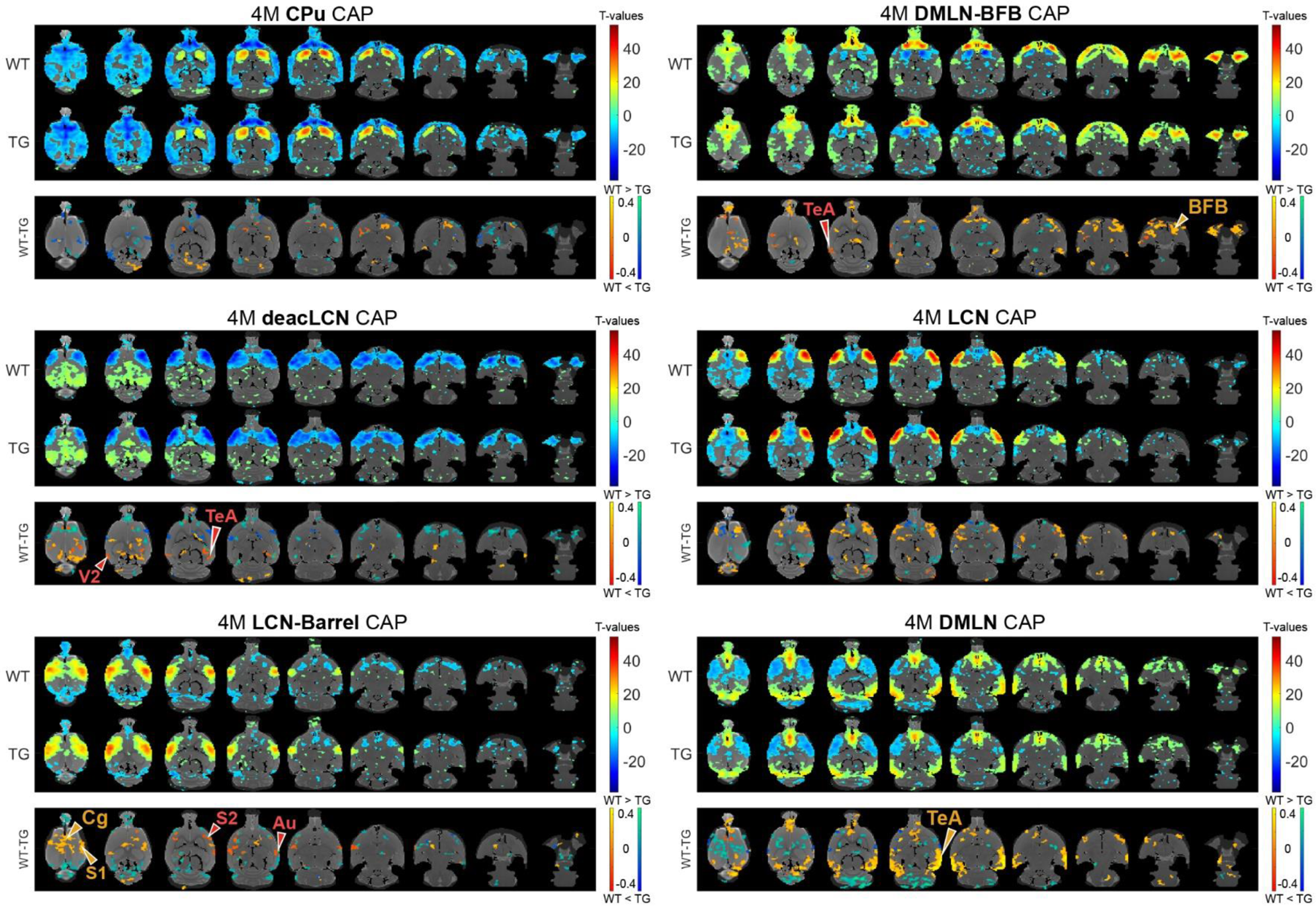
Genotypic group-level CAPs and post-hoc inter-group comparisons at the pre-plaque stage (4M). For each CAP, the **top panel** shows significantly activated and deactivated voxels represented as one-sample T-maps for each genotype. Voxels with positive T-values (yellow to red) indicate a significantly greater average activation than zero, whereas voxels with negative T-values (cyan to blue) indicate significantly greater deactivation than zero (p < 0.01; Bonferroni corrected). The **lower panels** (WT-TG) display the post-hoc comparisons (after 2-way ANOVA) of spatial activations between the two genotypes of voxels indicating a significant (FDR corrected, p < 0.05) difference in the (de)activation between the WT and TG CAPs. These post-hoc comparisons are made for voxels showing significant interaction effect among the voxels that show significant activation or deactivation in at least one group across all CAPs. The colour bars, ranging from red to yellow and blue to green, represent voxels that are respectively co-activated and co-deactivated within the WT group. Consequently, positive T-statistic values (in yellow and green) correspond to a significantly lower magnitude of activation in the TG group in contrast to the WT group, while negative T-statistic values (in red and blue) indicate a higher magnitude of activation in the TG group compared to the WT group. TeA, temporal association cortex; BFB, basal forebrain; S1, primary somatosensory cortex; S2, secondary somatosensory cortex; V2, secondary visual cortex; Cg, cingulate cortex; Au, auditory cortex.

**Figure 3.**
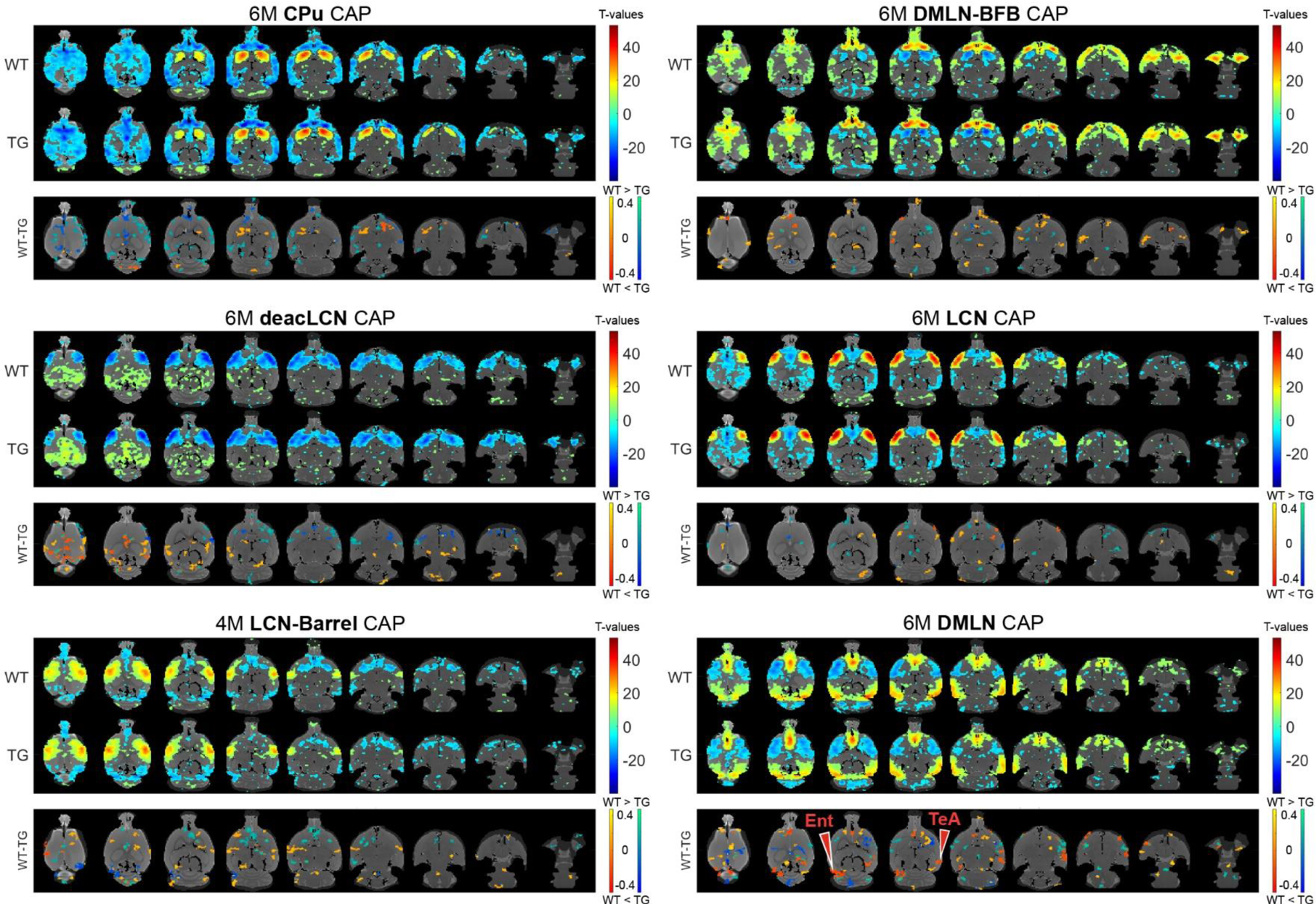
Group-level CAPs and comparisons at the early-plaque stage (6M). For each CAP, the **top panel** shows significantly activated and deactivated voxels represented as one-sample T-maps for each genotype. Voxels with positive T-values (yellow to red) indicate a significantly greater average activation than zero, whereas voxels with negative T-values (cyan to blue) indicate significantly greater deactivation than zero (p < 0.01; Bonferroni corrected). The **lower panels** (WT-TG) display the post-hoc comparisons (after 2-way ANOVA) of spatial activations between the two genotypes of voxels indicating a significant (FDR corrected, p < 0.05) difference in the (de)activation between the WT and TG CAPs. These post-hoc comparisons are made for voxels showing significant interaction effect among the voxels that show significant activation or deactivation in at least one group across all CAPs. The colour bars, ranging from red to yellow and blue to green, represent voxels that are respectively co-activated and co-deactivated within the WT group. Consequently, positive T-statistic values (in yellow and green) correspond to a significantly lower magnitude of activation in the TG group in contrast to the WT group, while negative T-statistic values (in red and blue) indicate a higher magnitude of activation in the TG group compared to the WT group. TeA, temporal association cortex; Ent, entorhinal cortex.

In the 4M *LCN-Barrel* CAP, the difference maps indicated a significantly lowered co-activation of central cortical regions (cingulate, primary and secondary motor, primary somatosensory cortices) in TG animals compared with WT animals. Moreover, TG animals exhibited highly localised hyperactivation in secondary somatosensory and auditory regions. In the *DMLN* CAP, the TGs had a lowered co-activation in the frontal and temporal association cortices, a decreased co-deactivation of motor cortices and a localized hyperactivation of the cingulate cortex. The *deacLCN* CAP in TG animals showed hyperactivation in the temporal association cortex and secondary visual cortex, along with hyperactivation of the cingulate cortex (as seen in the *DMLN* CAP). Hyperactivation of the temporal association cortex in TGs could also be seen in the *DMLN-BFB* CAP, along with reduced co-activation of BFB regions.

At 6M, most of the differences in spatial activation present at 4M were diminished (Figure 3). Notably, the pronounced lowered co-activation in TG animals across the BFB at 4M in the *DMLN-BFB* CAP was no longer observed at 6M, indicating an early transient effect. In the *DMLN* CAP, we observed localised hyperactivations in TG animals in the temporal association and entorhinal cortices.

### Temporal properties change in the *DMLN* CAP

To identify changes in the dynamics of brain states in disease vs control animals, we computed CAP occurrence percentage and duration across both groups at both ages. The occurrence percentage and duration of most CAPs was similar across groups and ages (Figure 4). The mean duration of the *DMLN* CAP significantly increased from 4M to 6M across both groups (Figure 4A). Interestingly, the *DMLN* CAP exhibited a group-dependent age effect in its occurrence percentage. The occurrence percentage of the *DMLN* CAP significantly increased only in the WT animals from 4M to 6M, but not in the TG animals that resulted in significant inter-genotypic difference at only 6M (Figure 4B).

**Figure 4.**
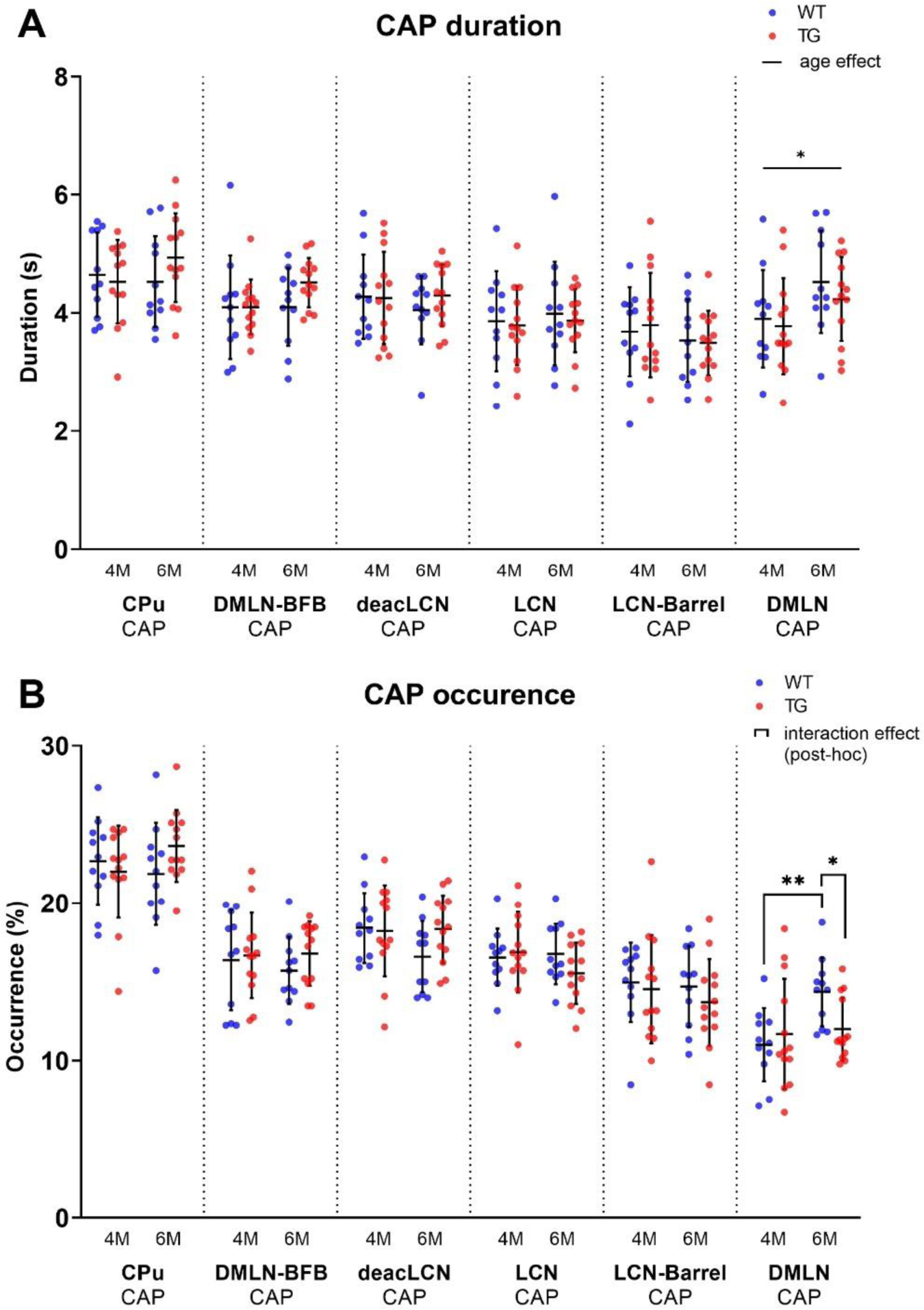
Temporal properties per age and genotype of all CAPs. Scatterplots of subject values per CAP, with the middle line indicating the mean and error bars indicating the SD. **(A)** Group-level CAP duration. Solid unbracketed line indicates age effect in the DMLN CAP (two-way RM ANOVA). **(B)** Group-level CAP occurrence percentage. Solid bracketed lines indicate significant multiple comparisons test (post-hoc two-way RM ANOVA, FDR corrected).

### Classification reaches higher-than-chance-level accuracy using CAP spatial features only

To assess the potential of using CAPs as a diagnostic biomarker, we explored the ability of CAPs to differentiate among the four combinations of genotype-age groups by using a predictive classifier. We utilized properties from the training subjects’ CAPs, derived from the complete image-series, as the classifier’s training features. Figure 5A displays the average classification accuracy when using temporal and spatial properties of CAPs, compared with the mean accuracy expected by random chance. In 4-class classification, temporal features (occurrence percentage and durations) of CAPs did not surpass random-chance accuracy, while classification using spatial features yielded significantly increased accuracy compared to the chance level (p = 3.98×10^−7^). The confusion matrix (Figure 5D) represents the classifier’s performance in predicting the correct labels for each genotype-age-group. With 4-class classification, the TG 4M group was most accurately predicted among the different subgroups, with an accuracy of 54%. Misclassification mostly occurred within the age group, between the genotypes. At 6M, the largest proportion of TG animals was accurately predicted (45%), with most misclassification occurring within the genotype but across age (25% misclassified as TG 4M). We additionally used a 2-class classification model per age where only genotypes were being predicted. Two-class classification using spatial features was significantly more accurate than the chance level at 4M (p = 0.0002; Figure 5B) but not at 6M (Figure 5C). At both ages, classification using temporal features did not surpass random chance levels. Highest prediction accuracy was achieved for the TG 4M subgroup, indicating its spatial properties were the most distinct from the other groups (Figure 5E).

**Figure 5.**
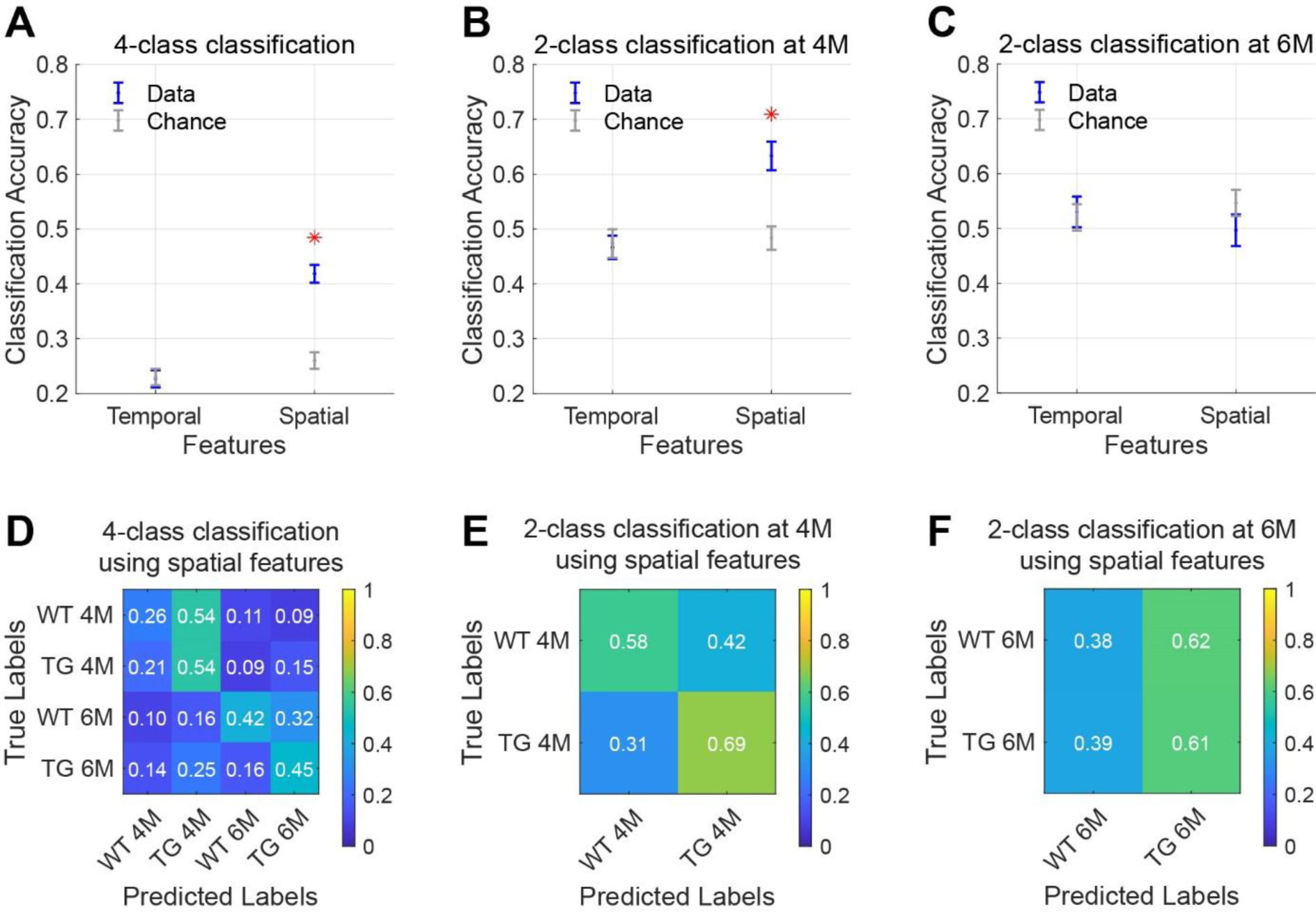
Classification of age and genotype groups using CAP features. **(A-C)** Accuracy of classification using temporal and spatial features in a 4-class model (**A**; WT 4M, TG 4M, WT 6M, TG 6M) or 2-class model per age (**B, C**; WT, TG). Data is in blue, chance-level accuracy in grey (mean ± SEM). Red asterisk indicates significantly higher mean accuracy compared to the chance level (FDR corrected, p < 0.05. **(D-F)** Confusion matrices representing performance of the classification model when using spatial features. Each value in the matrix indicates the proportion of test subjects that belong the group along the rows (true labels) and were predicted as the group on the columns (predicted labels), with the diagonal elements being correct predictions.

### Transition probabilities reveal subtle changes in network-state dynamics

Next, we investigated whether we could identify genotype-dependent changes in inter-CAP transitions, to gain insights into early changes in the dynamic organisation of intrinsic brain activity. To illustrate the pathways of CAP transitions at both 4M and 6M for WT and TG groups, we generated and graphically portrayed the transition probability matrices in each genotypic group at each age (Figure 6).

**Figure 6.**
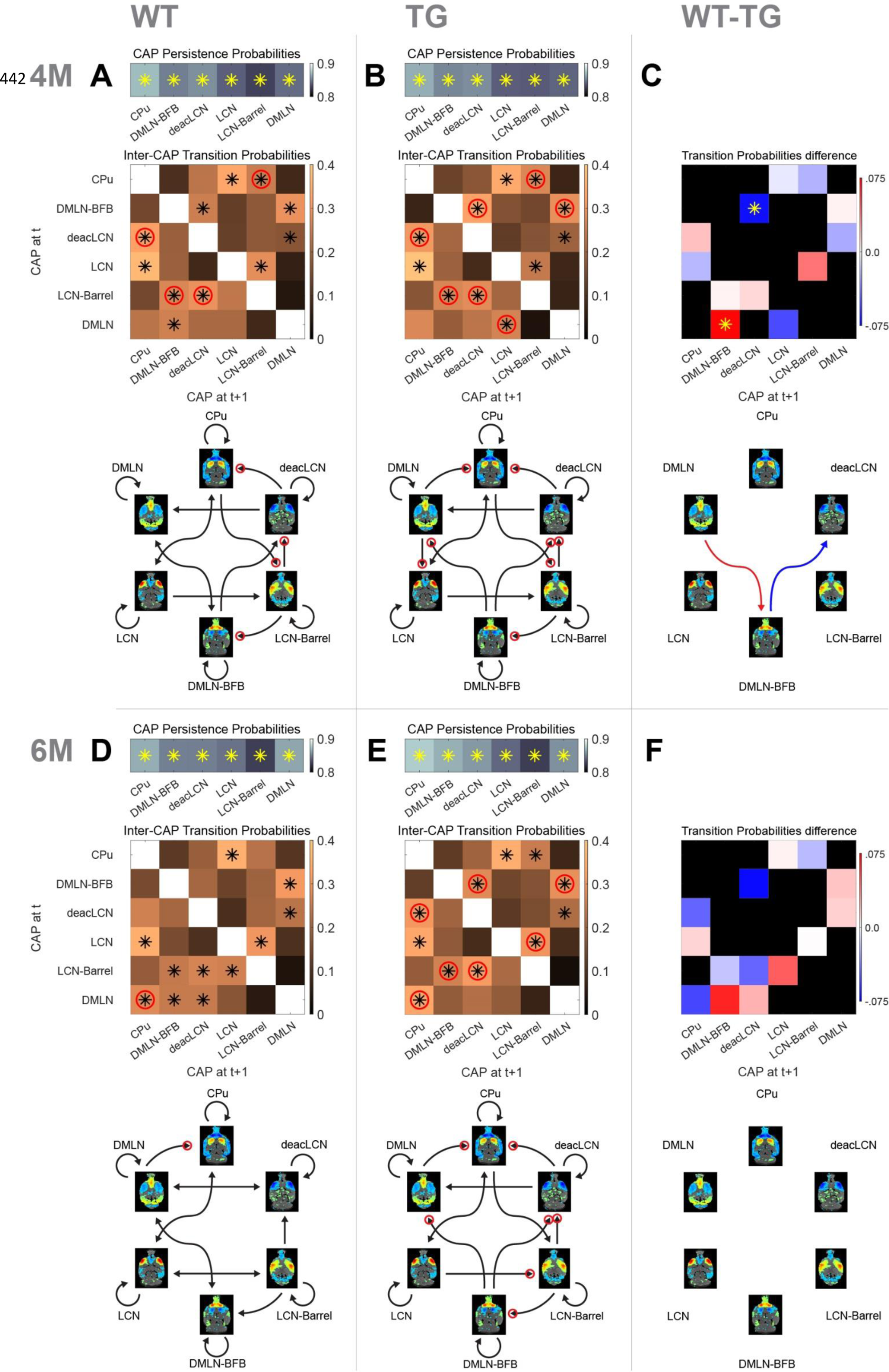
Probability matrices and schematic representations of transitions between CAPs. **(A, B, D, E)** Top row represents probabilities of persistent (self) transitions. Below that, inter-cap transition probabilities are represented in matrices. Colours indicate probability value, asterisk indicates transitions occurring significantly more frequently than the chance level, red circle indicates significantly higher probability of the transition in that direction compared to the opposite direction. Bottom schematic qualitatively represents significant transitions only. Each arrowhead corresponds to a significant transition in the probability matrix (asterisk), with the same red circle representation for directional preference. **(C, F)** Difference matrices between transition probabilities at 4M and 6M, respectively. Red colours indicate reduced transition probability in TG animals, while blue colours indicate increased transition probabilities. Asterisk indicates significant difference (p < 0.05, 2s T-test, FDR corrected). Schematic representation below shows significant differences only with red arrow indicating higher probability in WT while blue showing higher probability in TG.

We initially determined persistence probabilities (Figure 6A, B, D, E; top row), to examine the likelihood of CAPs transitioning into themselves, i.e., reoccurring in consecutive frames. A significant percentage of all transitions in all CAPs was persistent (∼85%), with each of the persistent transitions occurring significantly higher than the chance level. We observed mostly consistent persistence probabilities across the genotype-age-groups. The persistence probability of the *DMLN* CAP does increase from 4M to 6M in the WTs, but not in the TGs, tying in with the previously described differences in occurrence and duration of that CAP. Next, the transition probabilities from each CAP to other CAPs were calculated by considering only its non-persistent transitions, and visually represented in matrices (Figure 6A,B,D,E; matrices, off-diagonal).

At 4M, both genotypes exhibited several pairs of CAPs with significant transition probabilities (Figure 6A, B, black asterisks in matrices). Out of the 30 potential transition pathways, 11 were significantly more frequent than expected by the chance level in both genotypes (tested against distribution of surrogate transition probabilities per CAP-pair in permutation test, p < 0.05, FDR corrected). Notably, transitions from the *DMLN-BFB* and *deacLCN* CAPs to the *DMLN* CAP occurred with significantly higher probability than chance (p<0.05, FDR corrected) in both genotypes. In the WT animals, the *DMLN* CAP predominantly transitioned to the *DMLN-BFB* CAP, with this being the only significant outgoing transition (Figure 6A). In contrast, the TG animals displayed an alteration, with the primary significant outgoing transition from the *DMLN* CAP directed towards the *LCN* CAP (Figure 6B). We statistically compared transition probabilities, to uncover potential disease-induced changes in network dynamics. Transition probability from the *DMLN-BFB* CAP into the *deacLCN* CAP was significantly higher in TGs, and transitions from the *DMLN* CAP into the *DMLN-BFB* CAP were significantly less common than in the WTs (Figure 6C, p < 0.05, FDR corrected).

At 6M, transitions into the *DMLN* CAP remained significant from *DMLN-BFB* and *deacLCN* CAPs across both genotypes (Figure 6D, E, black asterisks in matrices). In the TG animals, significance of transitions remained largely the same as at 4M, except for the previously TG-specific significant transition of the *DMLN* CAP into the *LCN* CAP which was no longer significant. While transitions into the *DMLN* CAP did show the same consistent origin from *DMLN-BFB* and *deacLCN* CAP across genotypes and ages, in the TGs at 6M, *DMLN* CAP to *CPu* CAP is the only significant transition, compared to the more varied outgoing transitions from the *DMLN* CAP in the WTs. When directly comparing transition probabilities between the genotypes by permutation testing, no differences were statistically significant (p > 0.05; FDR corrected, Figure 6F).

### Directionality preference of transitions indicates genotypic difference at 6M

We also compared the probability of each transition p_ij_ with the other direction p_ji_ in each genotype-age-group, and tested the difference for significance, providing an indication of directional preference when transitioning between states. Through this analysis, we observed an additional distinct transition pattern at both ages, with a pronounced directionality preference for transitions originating from the *DMLN-BFB* CAP in the TG animals that was absent in WTs (Figure 6A,B,D,E; red circles on *DMLN-BFB* rows/arrows).

Overall, there was an increased directionality preference in the TG animals, with 7 (TG) vs 4 (WT) pairs showing directional preference (Figure 6A, B, red circles) at 4M. Directional preference for transitions *deacLCN*→*CPu, CPu*→*LCN-Barrel*, *LCN-Barrel*→*DMLN-BFB* and *LCN-Barrel*→*deacLCN* remained present across both genotypes, but the TG animals additionally showed increased directionality preference in transitions *DMLN-BFB*→*deacLCN, DMLN-BFB*→*DMLN* and *DMLN*→*LCN* (tested against distribution of surrogate transition probability differences p_ij_ – p_ji_ per CAP pair (I, j), p < 0.05, FDR corrected).

At 6M (Figure 6D, E), a specific directional transition from the *DMLN* into the *CPu* CAP was now apparent in both genotypes. Transitions in WTs now occurred mostly bidirectional with a nonsignificant directionality preference, with the sole exception of *DMLN*→*CPu* transition. In contrast, the transitions in TG animals still exhibited a directionality preference for 7 pairs of CAPs, 5 of which were identical to those at 4M.

## Discussion

In this study, we used resting-state CAPs to longitudinally examine properties of spontaneous whole-brain activity states in the TgF344-AD rat model of AD at the pre- and early-plaque stages of disease progression. We found six CAPs were the optimal number of states in this dataset, distinctly organised into CAP-anti-CAP pairs. DMLN activation was found in two of the CAPs, with one of them showing simultaneous BFB activation. In this *DMLN-BFB* CAP, BFB activation was reduced in the TG group only at the pre-plaque stage. Inter-genotypic differences in spatial activations of CAPs were diminished at the early-plaque stage. Analysis of CAP temporal properties revealed a reduction in the occurrence percentage of the *DMLN* CAP in the TG animals at the early-plaque stage. Cross-validated four-class classification of the animals into genotype-age-groups using spatial features of CAPs displayed significantly higher accuracy than chance. However, larger confusion between genotypes was found at the early-plaque stage which was confirmed when two genotypic groups were classified at each age; accuracy was significantly higher than the chance level at only the pre-plaque stage. Through characterisation of network state transitions, we identified genotypic differences in transition probabilities between two CAP pairs at the pre-plaque stage. TG animals showed increased probability for *DMLN*→*DMLN-BFB* transitions, and reduced probability for *DMLN-BFB*→*deacLCN* transitions. Interestingly, while several inter-CAP transitions at the pre-plaque stage exhibited directional preference in both genotypes, at the early-plaque stage, this directionality was observed only in the TG animals. In WT animals at the early-plaque stage, inter-CAP transitions were equally probable in both directions in most cases.

Our results, similar to earlier findings using CAP analysis in rodents (Gutierrez-Barragan et al., 2022, Adhikari et al., 2023), indicate a partition of 6 clusters as an optimal number to explain variance between the clusters, with diminishing returns using larger cluster numbers. Figure 1 These 6 clusters are distinctly organised into CAP-anti-CAP pairs. Across multiple CAPs, the DMLN and LCN co-activation patterns emerge as recognisable networks, previously described in the context of AD (Ibrahim et al., 2021, Liang et al., 2021, van den Berg et al., 2022). These networks are the respective rodent analogues of the DMN and TPN in humans. The DMN especially, involved in memory consolidation tasks, is the most prominent resting-state network, with AD patients typically exhibiting disrupted within-DMN connectivity (Grieder et al., 2018).

The *DMLN* CAP exhibits strong co-deactivation throughout cortical LCN regions (somatosensory, motor, and auditory cortex), thus further characterising it as a brain state resembling DMLN activity. Interestingly, it anticorrelates most strongly with the *LCN-Barrel* CAP, rather than the main *LCN* CAP itself. In the *DMLN-BFB* CAP, strong co-deactivation is strictly localised to the caudate putamen, extending slightly into the primary somatosensory cortex. It’s anti-correlated counterpart, *CPu* CAP, demonstrates highly localised co-activation in the caudate putamen only, with widespread cortical co-deactivation. Interestingly, the *DMLN-BFB* CAP shows the strongest co-activation in orbital and frontal association cortex together with BFB nuclei, while the *DMLN* CAP’s co-activation is more centred around the cingulate and prelimbic cortex. The separated DMLN-like CAPs we observe (*DMLN-BFB* and *DMLN*) are an example of how the single-frame resolution of CAPs provides enhanced statistical power, enabling a deeper and more granular understanding of the intricate and transient nature of brain connectivity. Another study comparing CAPs in awake and anesthetised mice (Gutierrez-Barragan et al., 2022) also found CAPs resembling DMN and LCN. Additionally, in their study, two CAPs showed high basal forebrain co-activation, one with hypothalamus and the other with auditory cortex, visual cortex, and amygdala. They also showed a CAP with high CPu co-activation, which was not isolated but co-activated mainly with cingulate, retrosplenial and prefrontal cortices. Their findings were consistent using either halothane or isoflurane-medetomidine anaesthesia.

One of the main outcomes of our spatial analysis of CAPs is the reduced co-activation in the BFB in TG animals at the pre-plaque stage, situated mainly in the *DMLN-BFB* CAP, confirming our earlier findings using QPPs (Geula et al., 2021). Basal forebrain cholinergic neurons (BFCN) experience early and substantial degeneration in AD, with the major neurodegenerative correlate being accumulation of pTau in neurofibrillary tangles (Geula et al., 2021). Consequently, BFCN degeneration is regarded as one of the hallmarks of AD (Shekari and Fahnestock, 2021). (Chen and Mobley, 2019)with evidence of decreased BFB volume before Aβ cortical spreading (Lozano-Montes et al., 2020, Nair et al., 2018). Given its implicated roles in modulation of cognition (through projections from the diagonal band of Broca, substantia innominata, and the nucleus basalis of Meynert to the cortex) (Espinosa et al., 2019, Li et al., 2015, Turchi et al., 2018) and its modulating function on brain activity through control on the prefrontal cortex (Espinosa et al., 2019, Li et al., 2015, Turchi et al., 2018), disruptions in BFB signalling caused by AD can lead to early disturbances in the brain’s network activities.

The differences in pre-plaque BFB co-activations we observe in the current study are like what was earlier described with quasi-periodic patterns (QPPs) in the same dataset (van den Berg et al., 2022). In that study, two short QPPs, called the DMLN and LCN QPPs consisting of coactivation of DMLN and LCN regions respectively, were found to be prominent. Co-activation of the BFB and the DMLN and the level of BFB activation was found be reduced at the pre-plaque stage in the TG animals. This reduction was not observed in the DMLN QPPs at the early-plaque stage. In the present study, while the *DMLN* and *DMLN-BFB* CAPs exhibit co-activation in similar hub regions involved in the DMLN (i.e., cingulate gyrus, retrosplenial cortex, orbitofrontal cortex, infralimbic cortex, visual cortex, and auditory cortex), the BFB activation was selective in the *DMLN-BFB* CAP (Figure 1A). Hence, the observed reduction of BFB co-activation in TG animals notably occurred mainly in the specific *DMLN-BFB* CAP, but not in the *DMLN* CAP, which is overall most closely resembling the DMLN (Figure 2 and Figure 3). In line with the earlier findings, these differences are pronounced during the pre-plaque stage, and mostly diminished at the early-plaque stage. QPPs are formed by averaging across recurring segments of scans of specific window size. As CAPs cover every frame, the overlap between QPP segments/occurrence and CAP occurrences can be calculated. We speculate that the previously described DMLN QPP involves brain states corresponding to both *DMLN-BFB* and *DMLN* CAPs, where the overlap of DMLN QPP segments with *DMLN-BFB* CAP would be significantly higher than that with *DMLN* CAP, especially at the pre-plaque stage.

In the *LCN-Barrel* CAP, anti-correlated to *DMLN* CAP and strongly resembling the LCN, we observed highly localised hyperactivations in somatosensory, auditory, and visual cortex regions in the TG animals. These are indicative of increased activation in the LCN, again corroborating previous findings (van den Berg et al., 2022). Like the other spatial differences, genotypic differences were most pronounced at the pre-plaque stage and diminished at the early-plaque stage.

Temporal properties of CAPs showed few differences between the genotypes, apart from the *DMLN* CAP. While mean duration of the *DMLN* CAP increased from the pre- to the early-plaque stage across both genotypes (Figure 4A), the occurrence percentage of the CAP increased only in the WTs, but not in the TG animals (Figure 4B). This suggests a healthy ageing effect on the occurrence of the *DMLN* CAP which is disrupted in the AD animals. In the zQ175 DN mouse model of Huntington’s disease, recent work employing a similar methodology (Adhikari et al., 2023) showed reduced durations of two anticorrelated CAPs, characterized by simultaneous co-activation of DMLN and co-deactivation of LCN and vice-versa, compared to the wild-type mice. In this work, multiple timepoints were investigated with only the final (10M) timepoint indicating significant differences in temporal metrics. In this light, we speculate that temporal metrics of the CAPs will be subject to stronger genotypic effects of AD at later ages, potentially involving other CAPs. Support for this hypothesis can be found in another study (Ma et al., 2020), where CAPs were studied in healthy elderly participants, patients with mild cognitive injury (MCI), and AD patients, showing that average dwell time in the DMN-like CAP showed a decreasing trend in AD.

Upon finding genotypic differences in both spatial and temporal metrics of the CAPs, we aimed to investigate their predictive power in identifying the genotypic status of the animals. In line with the earlier findings using this approach in Huntington’s disease mice (Adhikari et al., 2023), temporal properties of the CAPs did not perform classification better than chance in the current study. In contrast, classification using the spatial properties resulted in a significantly more accurate prediction of the genotype compared to the chance level. Interestingly, when stratifying the classifier per age, we found that accuracy of prediction reaches 65% using spatial features of the CAPs at the pre-plaque stage, but no significant improvement over the chance level at the early-plaque stage. Since earlier research using CAP analysis in AD mice at late-stage of AD development (18M) demonstrated near-perfect classification accuracy using CAP spatial features (Adhikari et al., 2020), we speculate that also in the TgF344-AD rat model, accuracy will further increase at later ages. We suspect accuracy of 4-class classification is lowered due to our timepoints being close (4M and 6M), combined with the earlier observation that genotypic spatial differences were less pronounced at the early-plaque than at pre-plaque stage.

One of the key advantages of CAP analysis over other dynamic FC methods is the additional dimension to characterisation of the dynamics, by not only revealing transient brain states and investigating their spatial and temporal properties, but by also characterising and quantifying the transitions between them. One finding of our CAP transition analysis is the consistent role of the *DMLN* CAP as distinct attractor for both the *DMLN-BFB* and *deacLCN* CAPs, across both genotypes and at both ages. These transitions do not only show significant probabilities; other transitions into the *DMLN* CAP have near-zero probabilities in each of the subgroups. Interpretation of this finding is two-fold. First, tying in with the earlier characterisation of the DMLN-like CAPs (*DMLN* and *DMLN-BFB)*, involvement of the BFB with the DMLN seems to be tied to a specific brain state that is separate from pure DMLN activity alone, but both states have a high affinity towards each other. Second, pure DMLN activation state rarely occurs directly out of LCN activity, but rather through more intermediate states; either the pronounced deactivation of the LCN (*deacLCN*) or the DMLN state with high involvement of the BFB (*DMLN-BFB*).

At the pre-plaque stage, the transition from *DMLN* to *DMLN-BFB* occurs significantly in the WTs (Figure 6A) but not in the TG animals (Figure 6B), in which its probability is significantly lower (Figure 6C). Moreover, while *DMLN*→*DMLN-BFB* is the only significant outgoing transition from the *DMLN* in pre-plaque WTs, the main outgoing transition from *DMLN* in TGs is directly to the *LCN*. This builds on observations of altered DMLN FC in AD patients (Greicius et al., 2004, Wang et al., 2013) and reduced segregation between DMN and TPN (Belloy et al., 2018b), with the current CAP analysis indicating an AD-induced shift in the flow of brain states involving the DMLN already at the pre-plaque stage. Additionally, we observe an increased probability of the *DMLN-BFB*→*deacLCN* transition in the TG animals.

At the early-plaque stage, transition probabilities are no longer different between the genotypes. Here, however, we observe a striking difference in the directionality preference of transitions. In the WT animals, besides the *DMLN*→*CPu* transition (1/11), none of the significant transitions exhibit directionality preferences. The TG animals exhibit directionality preference in 7 out of their 11 significant transitions. This suggests an altered rigidity of brain state flows, where TG animals seemingly exhibit more rigid transition patterns compared to the more flexible flow of brain states in the WTs. A modelling study, based on activity flow over altered intrinsic functional connections in AD, has previously reliably predicted task activations and related dysfunction in individuals at risk for AD (Mill et al., 2020). They proposed a mechanism where alterations in FC associated with AD would disrupt the flow of activations between regions, in turn leading to aberrant task activations. Our findings support the hypothesis that the resting-state flow of activations is altered, already at the very early stages of AD.

Comparing the current work to a previous study investigating persistence and transition probabilities in awake and anesthetised mice (Gutierrez-Barragan et al., 2022), we observe a few differences with our findings, which we can partly explain by the higher percentage of persistent transitions we detect. We observe that about 85% of our transitions are persistent, reflected in the longer durations of our CAPs while Gutierrez-Barragan et al. (Gutierrez-Barragan et al., 2022) find that about 60% of transitions are persistent. They also find a much larger variability in the values for persistence probabilities across CAPs compared to our study. Like our earlier findings and those of other groups (van den Berg et al., 2022, Tudela et al., 2019, Anckaerts et al., 2019), most of the spatial and temporal differences we can detect are mostly present at the pre-plaque stage but diminished in the early-plaque stage. This overall trend remains true for the transition probabilities; however, we can further nuance this finding with the observed alterations in directionality preference at the early-plaque stage. This nuanced characterisation of brain connectivity dynamics, inherent to CAP transition analysis, elucidates new insights into early alterations of intrinsic brain connectivity in AD.

Our study, while providing interesting insights, has several limitations. Our analysis was confined to male subjects only, which may limit the generalisability of our findings across genders. We used males only to limit variability and the number of animals required, however, sex differences in disease progression have been shown in humans (Webber et al., 2005) and AD models (Mielke, 2018), along with differences in pathological severity and behavioural deficits (Chaudry et al., 2022). We scanned under isoflurane and medetomidine anaesthesia, which is known to reduce FC in in subcortical areas. Previous research indicates that FC under this anaesthesia protocol closely resembles the awake state (Paasonen et al., 2018) and results in the most specific FC data compared to other protocols (Grandjean et al., 2023). However, a recent study comparing CAPs in awake and anesthetised mice brains has revealed significant alterations in topography of coactivations, their occurrences and temporal trajectories (Gutierrez-Barragan et al., 2022). Additionally, while our findings provide important insights into early-stage AD, the absence of direct behavioural outcome measures limits our ability to connect these neural changes to specific cognitive or behavioural symptoms. Finally, the use of BOLD signal as indirect measure of neural activity does not capture the entire scope and interplay of neural and hemodynamic alterations in AD. These limitations underscore the need for ongoing studies to further elucidate the complex relationship between hemodynamic, neural changes, and behavioural outcomes in AD pathology.

In conclusion, our study utilizing resting-state CAPs in the TgF344-AD rat model offers novel insights into altered brain functional dynamics at the early stages of Alzheimer’s disease (AD). We have validated key differences in the activation patterns and transitions of brain networks, particularly the DMLN and its interaction with the BFB, which are indicative of early disruptions in network dynamics associated with AD. Our findings confirm that these disruptions are most pronounced during the pre-plaque stage and tend to diminish as the disease progresses to the early-plaque stage. The *DMLN-BFB* and *DMLN* CAPs provide a deeper understanding of the interplay between different brain regions and their role in disease progression. Furthermore, our results underscore the potential of CAP analysis in predicting genotypic status based on spatial features, especially in early disease stages. This study not only corroborates previous findings on altered functional connectivity in AD but also advances our understanding of the nuanced changes in brain state transitions, offering a more detailed picture of the neurodegenerative process at the onset of plaque formation. Our approach, therefore, holds promise for early detection and better understanding of functional impact of the pathophysiology of AD, that may be relevant to assess the effects of potential therapeutic/disease-modifying interventions.

## References

Adhikari, M. H., Belloy, M. E., Van Der Linden, A., Keliris, G. A. & Verhoye, M. 2020. Resting-State Co-activation Patterns as Promising Candidates for Prediction of Alzheimer’s Disease in Aged Mice. Front Neural Circuits, 14, 612529.

Adhikari, M. H., Vasilkovska, T., Cachope, R., Tang, H., Liu, L., Keliris, G. A., Munoz-Sanjuan, I., Pustina, D., Van Der Linden, A. & Verhoye, M. 2023. Longitudinal investigation of changes in resting-state co-activation patterns and their predictive ability in the zQ175 DN mouse model of Huntington’s disease. Sci Rep, 13, 10194.

Agosta, F., Pievani, M., Geroldi, C., Copetti, M., Frisoni, G. B. & Filippi, M. 2012. Resting state fMRI in Alzheimer’s disease: beyond the default mode network. Neurobiol Aging, 33, 1564–78.

Alves, P. N., Foulon, C., Karolis, V., Bzdok, D., Margulies, D. S., Volle, E. & Thiebaut De Schotten, M. 2019. An improved neuroanatomical model of the default-mode network reconciles previous neuroimaging and neuropathological findings. Commun Biol, 2, 370.

An, Z., Tang, K., Xie, Y., Tong, C., Liu, J., Tao, Q., Consortium, D. & Feng, Y. 2024. Aberrant resting-state co-activation network dynamics in major depressive disorder. Transl Psychiatry, 14, 1.

Anckaerts, C., Blockx, I., Summer, P., Michael, J., Hamaide, J., Kreutzer, C., Boutin, H., Couillard-Despres, S., Verhoye, M. & Van Der Linden, A. 2019. Early functional connectivity deficits and progressive microstructural alterations in the TgF344-AD rat model of Alzheimer’s Disease: A longitudinal MRI study. Neurobiol Dis, 124, 93–107.

Badhwar, A., Tam, A., Dansereau, C., Orban, P., Hoffstaedter, F. & Bellec, P. 2017. Resting-state network dysfunction in Alzheimer’s disease: A systematic review and meta-analysis. Alzheimers Dement (Amst), 8, 73–85.

Belloy, M. E., Billings, J., Abbas, A., Kashyap, A., Pan, W. J., Hinz, R., Vanreusel, V., Van Audekerke, J., Van Der Linden, A., Keilholz, S. D., Verhoye, M. & Keliris, G. A. 2021. Resting Brain Fluctuations Are Intrinsically Coupled to Visual Response Dynamics. Cereb Cortex, 31, 1511–1522.

Belloy, M. E., Naeyaert, M., Abbas, A., Shah, D., Vanreusel, V., Van Audekerke, J., Keilholz, S. D., Keliris, G. A., Van Der Linden, A. & Verhoye, M. 2018a. Dynamic resting state fMRI analysis in mice reveals a set of Quasi-Periodic Patterns and illustrates their relationship with the global signal. Neuroimage, 180, 463–484.

Belloy, M. E., Shah, D., Abbas, A., Kashyap, A., Rossner, S., Van Der Linden, A., Keilholz, S. D., Keliris, G. A. & Verhoye, M. 2018b. Quasi-Periodic Patterns of Neural Activity improve Classification of Alzheimer’s Disease in Mice. Sci Rep, 8, 10024.

Berkowitz, L. E., Harvey, R. E., Drake, E., Thompson, S. M. & Clark, B. J. 2018. Progressive impairment of directional and spatially precise trajectories by TgF344-Alzheimer’s disease rats in the Morris Water Task. Sci Rep, 8, 16153.

Braak, H., Thal, D. R., Ghebremedhin, E. & Del Tredici, K. 2011. Stages of the pathologic process in Alzheimer disease: age categories from 1 to 100 years. J Neuropathol Exp Neurol, 70, 960–9.

Calhoun, V. D., Miller, R., Pearlson, G. & Adali, T. 2014. The chronnectome: time-varying connectivity networks as the next frontier in fMRI data discovery. Neuron, 84, 262–74.

Chaudry, O., Ndukwe, K., Xie, L., Figueiredo-Pereira, M., Serrano, P. & Rockwell, P. 2022. Females exhibit higher GluA2 levels and outperform males in active place avoidance despite increased amyloid plaques in TgF344-Alzheimer’s rats. Sci Rep, 12, 19129.

Chen, X. Q. & Mobley, W. C. 2019. Exploring the Pathogenesis of Alzheimer Disease in Basal Forebrain Cholinergic Neurons: Converging Insights From Alternative Hypotheses. Front Neurosci, 13, 446.

Cohen, R. M., Rezai-Zadeh, K., Weitz, T. M., Rentsendorj, A., Gate, D., Spivak, I., Bholat, Y., Vasilevko, V., Glabe, C. G., Breunig, J. J., Rakic, P., Davtyan, H., Agadjanyan, M. G., Kepe, V., Barrio, J. R., Bannykh, S., Szekely, C. A., Pechnick, R. N. & Town, T. 2013. A transgenic Alzheimer rat with plaques, tau pathology, behavioral impairment, oligomeric abeta, and frank neuronal loss. J Neurosci, 33, 6245–56.

Espinosa, N., Alonso, A., Morales, C., Espinosa, P., Chavez, A. E. & Fuentealba, P. 2019. Basal Forebrain Gating by Somatostatin Neurons Drives Prefrontal Cortical Activity. Cereb Cortex, 29, 42–53.

Geula, C., Dunlop, S. R., Ayala, I., Kawles, A. S., Flanagan, M. E., Gefen, T. & Mesulam, M. M. 2021. Basal forebrain cholinergic system in the dementias: Vulnerability, resilience, and resistance. J Neurochem, 158, 1394–1411.

Grandjean, J., Desrosiers-Gregoire, G., Anckaerts, C., Angeles-Valdez, D., Ayad, F., Barrière, D. A., Blockx, I., Bortel, A., Broadwater, M., Cardoso, B. M., Célestine, M., Chavez-Negrete, J. E., Choi, S., Christiaen, E., Clavijo, P., Colon-Perez, L., Cramer, S., Daniele, T., Dempsey, E., Diao, Y., Doelemeyer, A., Dopfel, D., Dvořáková, L., Falfán-Melgoza, C., Fernandes, F. F., Fowler, C. F., Fuentes-Ibañez, A., Garin, C. M., Gelderman, E., Golden, C. E. M., Guo, C. C. G., Henckens, M., Hennessy, L. A., Herman, P., Hofwijks, N., Horien, C., Ionescu, T. M., Jones, J., Kaesser, J., Kim, E., Lambers, H., Lazari, A., Lee, S. H., Lillywhite, A., Liu, Y., Liu, Y. Y., López-Castro, A., López-Gil, X., Ma, Z., Macnicol, E., Madularu, D., Mandino, F., Marciano, S., Mcauslan, M. J., Mccunn, P., Mcintosh, A., Meng, X., Meyer-Baese, L., Missault, S., Moro, F., Naessens, D. M. P., Nava-Gomez, L. J., Nonaka, H., Ortiz, J. J., Paasonen, J., Peeters, L. M., Pereira, M., Perez, P. D., Pompilus, M., Prior, M., Rakhmatullin, R., Reimann, H. M., Reinwald, J., Del Rio, R. T., Rivera-Olvera, A., Ruiz-Pérez, D., Russo, G., Rutten, T. J., Ryoke, R., Sack, M., Salvan, P., Sanganahalli, B. G., Schroeter, A., Seewoo, B. J., Selingue, E., Seuwen, A., Shi, B., Sirmpilatze, N., Smith, J. A. B., Smith, C., Sobczak, F., Stenroos, P. J., Straathof, M., Strobelt, S., Sumiyoshi, A., Takahashi, K., Torres-García, M. E., Tudela, R., Van Den Berg, M., Van Der Marel, K., et al. 2023. A consensus protocol for functional connectivity analysis in the rat brain. Nat Neurosci, 26, 673–681.

Greicius, M. D., Srivastava, G., Reiss, A. L. & Menon, V. 2004. Default-mode network activity distinguishes Alzheimer’s disease from healthy aging: evidence from functional Mri. Proc Natl Acad Sci U S A, 101, 4637–42.

Grieder, M., Wang, D. J. J., Dierks, T., Wahlund, L. O. & Jann, K. 2018. Default Mode Network Complexity and Cognitive Decline in Mild Alzheimer’s Disease. Front Neurosci, 12, 770.

Gutierrez-Barragan, D., Basson, M. A., Panzeri, S. & Gozzi, A. 2019. Infraslow State Fluctuations Govern Spontaneous fMRI Network Dynamics. Curr Biol, 29, 2295–2306 e5.

Gutierrez-Barragan, D., Singh, N. A., Alvino, F. G., Coletta, L., Rocchi, F., De Guzman, E., Galbusera, A., Uboldi, M., Panzeri, S. & Gozzi, A. 2022. Unique spatiotemporal fMRI dynamics in the awake mouse brain. Curr Biol, 32, 631–644 e6.

Hutchison, R. M., Womelsdorf, T., Allen, E. A., Bandettini, P. A., Calhoun, V. D., Corbetta, M., Della Penna, S., Duyn, J. H., Glover, G. H., Gonzalez-Castillo, J., Handwerker, D. A., Keilholz, S., Kiviniemi, V., Leopold, D. A., De Pasquale, F., Sporns, O., Walter, M. & Chang, C. 2013. Dynamic functional connectivity: promise, issues, and interpretations. Neuroimage, 80, 360–78.

Ibrahim, B., Suppiah, S., Ibrahim, N., Mohamad, M., Hassan, H. A., Nasser, N. S. & Saripan, M. I. 2021. Diagnostic power of resting-state fMRI for detection of network connectivity in Alzheimer’s disease and mild cognitive impairment: A systematic review. Hum Brain Mapp, 42, 2941–2968.

Joo, I. L., Lai, A. Y., Bazzigaluppi, P., Koletar, M. M., Dorr, A., Brown, M. E., Thomason, L. A., Sled, J. G., Mclaurin, J. & Stefanovic, B. 2017. Early neurovascular dysfunction in a transgenic rat model of Alzheimer’s disease. Sci Rep, 7, 46427.

Lee, K., Ji, J. L., Fonteneau, C., Berkovitch, L., Rahmati, M., Pan, L., Repovs, G., Krystal, J. H., Murray, J. D. & Anticevic, A. 2023. Human brain state dynamics reflect individual neuro-phenotypes. bioRxiv.

Li, M., Dahmani, L., Wang, D., Ren, J., Stocklein, S., Lin, Y., Luan, G., Zhang, Z., Lu, G., Galie, F., Han, Y., Pascual-Leone, A., Wang, M., Fox, M. D. & Liu, H. 2021. Co-activation patterns across multiple tasks reveal robust anti-correlated functional networks. Neuroimage, 227, 117680.

Li, W., Motelow, J. E., Zhan, Q., Hu, Y. C., Kim, R., Chen, W. C. & Blumenfeld, H. 2015. Cortical network switching: possible role of the lateral septum and cholinergic arousal. Brain Stimul, 8, 36–41.

Liang, L., Yuan, Y., Wei, Y., Yu, B., Mai, W., Duan, G., Nong, X., Li, C., Su, J., Zhao, L., Zhang, Z. & Deng, D. 2021. Recurrent and concurrent patterns of regional Bold dynamics and functional connectivity dynamics in cognitive decline. Alzheimers Res Ther, 13, 28.

Liu, X. & Duyn, J. H. 2013. Time-varying functional network information extracted from brief instances of spontaneous brain activity. Proc Natl Acad Sci U S A, 110, 4392–7.

Liu, X., Zhang, N., Chang, C. & Duyn, J. H. 2018. Co-activation patterns in resting-state fMRI signals. Neuroimage, 180, 485–494.

Lozano-Montes, L., Dimanico, M., Mazloum, R., Li, W., Nair, J., Kintscher, M., Schneggenburger, R., Harvey, M. & Rainer, G. 2020. Optogenetic Stimulation of Basal Forebrain Parvalbumin Neurons Activates the Default Mode Network and Associated Behaviors. Cell Rep, 33, 108359.

Lu, H., Zou, Q., Gu, H., Raichle, M. E., Stein, E. A. & Yang, Y. 2012. Rat brains also have a default mode network. Proc Natl Acad Sci U S A, 109, 3979–84.

Lurie, D. J., Kessler, D., Bassett, D. S., Betzel, R. F., Breakspear, M., Kheilholz, S., Kucyi, A., Liegeois, R., Lindquist, M. A., Mcintosh, A. R., Poldrack, R. A., Shine, J. M., Thompson, W. H., Bielczyk, N. Z., Douw, L., Kraft, D., Miller, R. L., Muthuraman, M., Pasquini, L., Razi, A., Vidaurre, D., Xie, H. & Calhoun, V. D. 2020. Questions and controversies in the study of time-varying functional connectivity in resting fMRI. Netw Neurosci, 4, 30–69.

Ma, X., Zhuo, Z., Wei, L., Ma, Z., Li, Z., Li, H. & ALZHEIMER’s Disease Neuroimaging, I. 2020. Altered Temporal Organization of Brief Spontaneous Brain Activities in Patients with Alzheimer’s Disease. Neuroscience, 425, 1–11.

Majeed, W., Magnuson, M., Hasenkamp, W., Schwarb, H., Schumacher, E. H., Barsalou, L. & Keilholz, S. D. 2011. Spatiotemporal dynamics of low frequency BOLD fluctuations in rats and humans. Neuroimage, 54, 1140–50.

Maltbie, E., Yousefi, B., Zhang, X., Kashyap, A. & Keilholz, S. 2022. Comparison of Resting-State Functional Mri Methods for Characterizing Brain Dynamics. Front Neural Circuits, 16, 681544.

Mielke, M. M. 2018. Sex and Gender Differences in Alzheimer’s Disease Dementia. Psychiatr Times, 35, 14–17.

Mill, R. D., Gordon, B. A., Balota, D. A. & Cole, M. W. 2020. Predicting dysfunctional age-related task activations from resting-state network alterations. Neuroimage, 221, 117167.

Munoz-Moreno, E., Tudela, R., Lopez-Gil, X. & Soria, G. 2018. Early brain connectivity alterations and cognitive impairment in a rat model of Alzheimer’s disease. Alzheimers Res Ther, 10, 16.

Munoz-Moreno, E., Tudela, R., Lopez-Gil, X. & Soria, G. 2020. Brain connectivity during Alzheimer’s disease progression and its cognitive impact in a transgenic rat model. Netw Neurosci, 4, 397–415.

Nair, J., Klaassen, A. L., Arato, J., Vyssotski, A. L., Harvey, M. & Rainer, G. 2018. Basal forebrain contributes to default mode network regulation. Proc Natl Acad Sci U S A, 115, 1352–1357.

Paasonen, J., Stenroos, P., Salo, R. A., Kiviniemi, V. & Gröhn, O. 2018. Functional connectivity under six anesthesia protocols and the awake condition in rat brain. Neuroimage, 172, 9–20.

Pentkowski, N. S., Berkowitz, L. E., Thompson, S. M., Drake, E. N., Olguin, C. R. & Clark, B. J. 2018. Anxiety-like behavior as an early endophenotype in the TgF344-AD rat model of Alzheimer’s disease. Neurobiol Aging, 61, 169–176.

Ramzan, F., Khan, M. U. G., Rehmat, A., Iqbal, S., Saba, T., Rehman, A. & Mehmood, Z. 2019. A Deep Learning Approach for Automated Diagnosis and Multi-Class Classification of Alzheimer’s Disease Stages Using Resting-State fMRI and Residual Neural Networks. J Med Syst, 44, 37.

Rorabaugh, J. M., Chalermpalanupap, T., Botz-Zapp, C. A., Fu, V. M., Lembeck, N. A., Cohen, R. M. & Weinshenker, D. 2017. Chemogenetic locus coeruleus activation restores reversal learning in a rat model of Alzheimer’s disease. Brain, 140, 3023–3038.

Shekari, A. & Fahnestock, M. 2021. Chapter 13 - Cholinergic neurodegeneration in Alzheimer disease mouse models. In: Swaab, D. F., Buijs, R. M., Kreier, F., Lucassen, P. J. & Salehi, A. (eds.) Handbook of Clinical Neurology. Elsevier.

Sheline, Y. I. & Raichle, M. E. 2013. Resting state functional connectivity in preclinical Alzheimer’s disease. Biol Psychiatry, 74, 340–7.

Tagliazucchi, E., Balenzuela, P., Fraiman, D. & Chialvo, D. R. 2012. Criticality in large-scale brain FMRI dynamics unveiled by a novel point process analysis. Front Physiol, 3, 15.

Tournier, B. B., Barca, C., Fall, A. B., Gloria, Y., Meyer, L., Ceyzeriat, K. & Millet, P. 2021. Spatial reference learning deficits in absence of dysfunctional working memory in the TgF344-AD rat model of Alzheimer’s disease. Genes Brain Behav, 20, e12712.

Tudela, R., Munoz-Moreno, E., Sala-Llonch, R., Lopez-Gil, X. & Soria, G. 2019. Resting State Networks in the TgF344-AD Rat Model of Alzheimer’s Disease Are Altered From Early Stages. Front Aging Neurosci, 11, 213.

Turchi, J., Chang, C., Ye, F. Q., Russ, B. E., Yu, D. K., Cortes, C. R., Monosov, I. E., Duyn, J. H. & Leopold, D. A. 2018. The Basal Forebrain Regulates Global Resting-State fMRI Fluctuations. Neuron, 97, 940–952e4.

Van Den Berg, M., Adhikari, M. H., Verschuuren, M., Pintelon, I., Vasilkovska, T., Van Audekerke, J., Missault, S., Heymans, L., Ponsaerts, P., De Vos, W. H., Van Der Linden, A., Keliris, G. A. & Verhoye, M. 2022. Altered basal forebrain function during whole-brain network activity at pre- and early-plaque stages of Alzheimer’s disease in TgF344-AD rats. Alzheimers Res Ther, 14, 148.

Van Den Heuvel, M. P. & Sporns, O. 2013. Network hubs in the human brain. Trends Cogn Sci, 17, 683–96.

Vasilkovska, T., Adhikari, M. H., Van Audekerke, J., Salajeghe, S., Pustina, D., Cachope, R., Tang, H., Liu, L., Munoz-Sanjuan, I., Van Der Linden, A. & Verhoye, M. 2023. Resting-state fMRI reveals longitudinal alterations in brain network connectivity in the zQ175dn mouse model of Huntington’s disease. Neurobiol Dis, 181, 106095.

Wang, Y., Risacher, S. L., West, J. D., Mcdonald, B. C., Magee, T. R., Farlow, M. R., Gao, S., O’neill, D. P. & Saykin, A. J. 2013. Altered default mode network connectivity in older adults with cognitive complaints and amnestic mild cognitive impairment. J Alzheimers Dis, 35, 751–60.

Webber, K. M., Casadesús, G., Perry, G., Atwood, C. S., Bowen, R. L. & Smith, M. A. 2005. Gender Differences in Alzheimer Disease. Alzheimer Disease & Associated Disorders.

Xu, N., Lagrow, T. J., Anumba, N., Lee, A., Zhang, X., Yousefi, B., Bassil, Y., Clavijo, G. P., Khalilzad Sharghi, V., Maltbie, E., Meyer-Baese, L., Nezafati, M., Pan, W. J. & Keilholz, S. 2022. Functional Connectivity of the Brain Across Rodents and Humans. Front Neurosci, 16, 816331.

Yousefi, B. & Keilholz, S. 2021. Propagating patterns of intrinsic activity along macroscale gradients coordinate functional connections across the whole brain. Neuroimage, 231, 117827.

Yousefi, B., Shin, J., Schumacher, E. H. & Keilholz, S. D. 2018. Quasi-periodic patterns of intrinsic brain activity in individuals and their relationship to global signal. Neuroimage, 167, 297–308.

